# New insights into the structure of Comirnaty Covid-19 vaccine: A theory on soft nanoparticles with mRNA-lipid supercoils stabilized by hydrogen bonds

**DOI:** 10.1101/2022.12.02.518611

**Authors:** János Szebeni, Bálint Kiss, Tamás Bozó, Keren Turjeman, Yael Levi-Kalisman, Yechezkel Barenholz, Miklós Kellermayer

## Abstract

Despite the worldwide success of mRNA-LNP Covid-19 vaccines, the nanoscale structure of these formulations is still poorly understood. To fill this gap, we used a combination of atomic force microscopy (AFM), dynamic light scattering (DLS), transmission electron microscopy (TEM), cryogenic transmission electron microscopy (cryo-TEM) and the determination of LNP pH gradient to analyze the nanoparticles (NPs) in BNT162b2 (Comirnaty), comparing it with the well characterized pegylated liposomal doxorubicin (Doxil). Comirnaty NPs had similar size to Doxil, however, unlike Doxil liposomes, wherein the stable ammonium and pH gradient enables accumulation of ^14^C-methylamine in the intraliposomal aqueous phase, Comirnaty LNPs lack such pH gradient in spite of the fact that the pH 4, at which LNPs are prepared, is raised to pH 7.2 after loading of the mRNA. Mechanical manipulation of Comirnaty NPs with AFM revealed soft, compliant structures. The sawtooth-like force transitions seen during cantilever retraction implies that molecular strands, corresponding to mRNA, can be pulled out of NPs, and the process is accompanied by stepwise rupture of mRNA-lipid bonds. Unlike Doxil, cryo-TEM of Comirnaty NPs revealed a granular, solid core enclosed by mono- and bilayers. Negative staining TEM shows 2-5 nm electron-dense spots in the liposom’s interior that are aligned into strings, semicircles, or labyrinth-like networks, which may imply crosslink-stabilized supercoils. The neutral intra-LNP core questions the dominance of ionic interactions holding together this scaffold, raising the alternative possibility of hydrogen bonding between the mRNA and the lipids. Such interaction, described previously for another mRNA/lipid complex, is consistent with the steric structure of ionizable lipid in Comirnaty, ALC-0315, displaying free =O and -OH groups. It is hypothesized that the latter groups can get into steric positions that enable hydrogen bonding with the nitrogenous bases in the mRNA. These newly recognized structural features of mRNA-LNP may be important for the vaccine’s efficacy.

## Introduction

The worldwide use of Pfizer/BioNTech’s Comirnaty (BNT162b2) and Moderna’s Spikevax (mRNA-1273) vaccines against Covid-19 brought substantial public and scientific interest and scrutiny. They represent a new approach of immunization based on mRNA-containing lipid nanoparticles (mRNA-LNPs), wherein the mRNA encodes the virus’s spike protein (S-protein). Following translation, the protein presents antigen for the immune system to develop specific immunity against the virus. This approach of vaccination stems from the success of LNPs in gene therapy and lipophilic drug delivery^1, 2^ that utilizes ionizable lipids displaying positive charge at low pH (IPC lipids), neutral PEGylated lipids, neutral membrane-forming phospholipids and cholesterol.^3–5^ However, the compact LNPs are very different from the clinically applied bilayer liposomes with clearly discernible internal aqueous compartment. Yet there is little information about the molecular buildup of fully filled LNPs in general, and Comirnaty in particular. Accordingly, the goal of the present study was to employ a combination of state-of-art nanostructure analysis techniques to better understand the structure of Comirnaty in comparison with Doxil, a well-characterized liposome control, whose chemical composition is compared to that of Comirnaty in Supplement Table 1.

We used atomic force microscopy (AFM) imaging and force spectroscopy to unveil the 3Dshape and nanomechanical properties of Comirnaty, dynamic light scattering (DLS) to ascertain the averaged hydrodynamic size of NPs, cryogenic-transmission electron microscopy (cryo-TEM) to image the individual LNPs, their size and shape and internal fine structure, and radiolabel equilibrium measurements to explore the presence of a pH gradient and permeability of the LNP coating layer to small molecules, and to measure intraparticle pH. Our data, taken together with a recent description of IPC lipid clusters binding to mRNA with hydrogen bonds^6^ led us to propose a new model of Comirnaty structure wherein the mRNA supercoils are stabilized by hydrogen bonds with IPC-lipids with weaker or absent ionic interactions, and intermittent mono- and bilayered membranes making up the coat.

## RESULTS AND DISCUSSION

### AFM images of Comirnaty as administered in people and after storage of expired samples

This study focuses on the structure of Comirnaty as administered to people, also noting changes observed after 1 day storage at 4°C, which is irrelevant regarding the human application of the vaccine but reveals information that help understanding the structure of these NPs.

Fig. 1 shows AFM images of Comirnaty and Doxil NPs under different conditions with regards to storage time and temperature. It is seen, particularly in the 3D reconstituted versions of images (lower panels), that the freshly diluted vaccine (Fig. 1A and E), representing the inoculum used for injection into humans, consists of monodisperse, spherical NPs of about 120-150 nm apparent diameter and 40-60 nm topographical height (54.6±19.3 nm, mean ± SD, n=148), corresponding to slightly flattened surface-adsorbed NPs. One-day storage of diluted vaccine at 4°C, which represents samples not recommended for human use, led to striking differences in sample morphology (Fig.1B and F). Namely, both the size and shape of NPs became heterogeneous, and large flat patches were visible on the supporting surface. Refreezing the stored samples, which, too, is excluded by the manufacturer for human use, led to even more heterogeneous, disperse NPs with substantial variety of fragment shapes in the ~5 to ~300 nm range (Figure 1C and G). The freshly opened Doxil (Fig. 1D and H) showed similar monodisperse, spherical NPs as are the freshly suspended Comirnaty NPs (Fig 1A and E) but their topographical height was lower (28.5±7.8 nm, mean ± SD).

**Figure 1.**
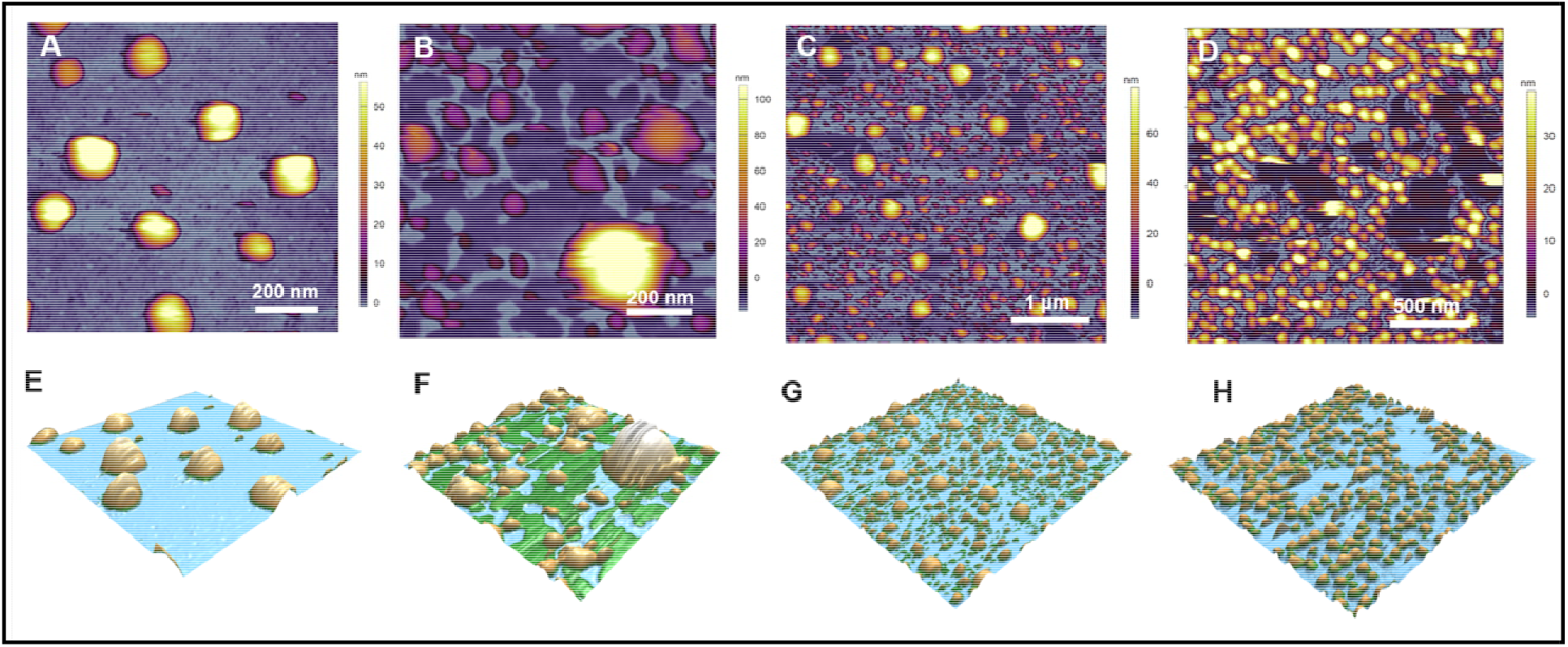
AFM height-contrast images (A-D) and corresponding 3D reconstructions (E-F) of Comirnaty vaccine and Doxil immobilized on glass surface. (A) and (E) show a freshly diluted Comirnaty sample representing the jab inoculated into the deltoid muscle; (B) and (F) show a 1-day old sample stored at 4°C, representing the unused leftover; (C) and (G) show a refrozen sample, representing unintended acceleration of fragmentation by refreezing the leftover vaccine; (D) and (H) show a freshly opened Doxil sample.

Regarding the morphological changes of Comirnaty upon storage, as shown by AFM, the increase of particle size without segmentation and losing of smooth surface may be explained by time-dependent association of NPs, most likely via fusion. The decrease of particle size, in turn, suggests that loss of phospholipids has occurred in some liposomes, giving rise of flat patches in the background that correspond to lipid layers stretched on the surface (Fig 1B, F).

### DLS analysis of the hydrodynamic size and size distribution of freshly diluted Comirnaty and Doxil

As shown in Table 1, the freshly diluted Comirnaty and Doxil NPs had essentially similar hydrodynamic size (diameter: 80-85 nm), but the homogeneity of Comirnaty was slightly lower than that of Doxil, as reflected in the higher span of size distribution and higher polydispersity index (Pdi) of Comirnaty NPs compared to Doxil. Also, the surface charge (Zp) of Doxil (−31 +/− 5.1 (mean +/− SD, n=3) was more negative than that of Comirnaty (8.6 +/− 5.3, mean +/− SD, n= 3), as determined in 1.5 mM sodium nitrate. Nevertheless, these data are consistent with the AFM images inasmuch as both preparations consist of spherical NPs of about the same size. As for the discrepancy between the hydrodynamic diameter measured by DLS and topographical height of NPs, measured by AFM, it can be attributed to the different focuses of the 2 methods; global estimate of NP size in solution vs. 3D shape of individual NPs adhered to a surface, respectively.

**Table 1.** DLS analysis of the hydrodynamic size and size distribution of NPs in freshly diluted Comirnaty and Doxil

| NPs | Z-Ave | D(v)<br>10 | D(v)<br>50 | D(v)<br>90 | D(i)<br>10 | D(i)<br>50 | D(i)<br>90 | SPAN<br>(i) | Pdi | Zp |
| --- | --- | --- | --- | --- | --- | --- | --- | --- | --- | --- |
|  |  | Batch A |  |  | Batch B |  |  |  |  |  |
| Comirnaty | 84.4<br>±0.3 | 40.6<br>±0.9 | 61.2<br>±0.9 | 109±<br>1.7 | 53<br>±0.3 | 88.7<br>±1 | 155.3<br>±0.6 | 1.1 | 0.20 | -8.6<br>±2.6 |
| Doxil | 81.5<br>±0.4 | 48.7<br>±1.0 | 68.9<br>±1.0 | 105±<br>0.6 | 58<br>±0.9 | 84.5<br>±0.3 | 123<br>±0.6 | 0.8 | 0.06 | -31<br>±5.1 |

Entries are mean ± SD, n=3 measurement in each of two different sample batches. Span is defined as $D_{90}-D_{10}/D_{50}$ where $D_{10}$ , $D_{50}$ and $D_{90}$ are the percentiles under the size distribution curve; PDI, polydispersity index. Further details are described in the Methods section.

### Nanomechanical properties of freshly diluted Comirnaty NPs

To explore the nanomechanical properties of vaccine NPs administered to humans, we analyzed the force-distance curves obtained in freshly diluted Comirnaty samples upon NP indentation with the AFM’s cantilever. Fig. 2A shows a representative force curve recorded during an indentation cycle. During tip approach (red curve), following a constant-force region reflecting the lack of load as the cantilever moved towards the NP surface, a rise in force was apparent (phase 1). The distance (cca. 50 nm) at which the force began to rise, corresponds well to topographical height of NPs, indicating the point of contact between the surface-adsorbed particle and the AFM tip. The apparent linear increase of force is a clear sign of elastic compression of the NP. The slope of the linear fit in this region provided a stiffness of ~9 pN/nm (Table 2), which is slightly smaller than that of dimyristoyl-phosphatidylcholine (DMPC) liposomes^7^ and an order of magnitude smaller than that of dipalmitoyl-phosphatidylcholine (DPPC) liposomes^8^ with roughly the same radius (i.e., liquid-disordered and liquid-ordered membrane liposomes at room temperature, respectively).

**Figure 2.**
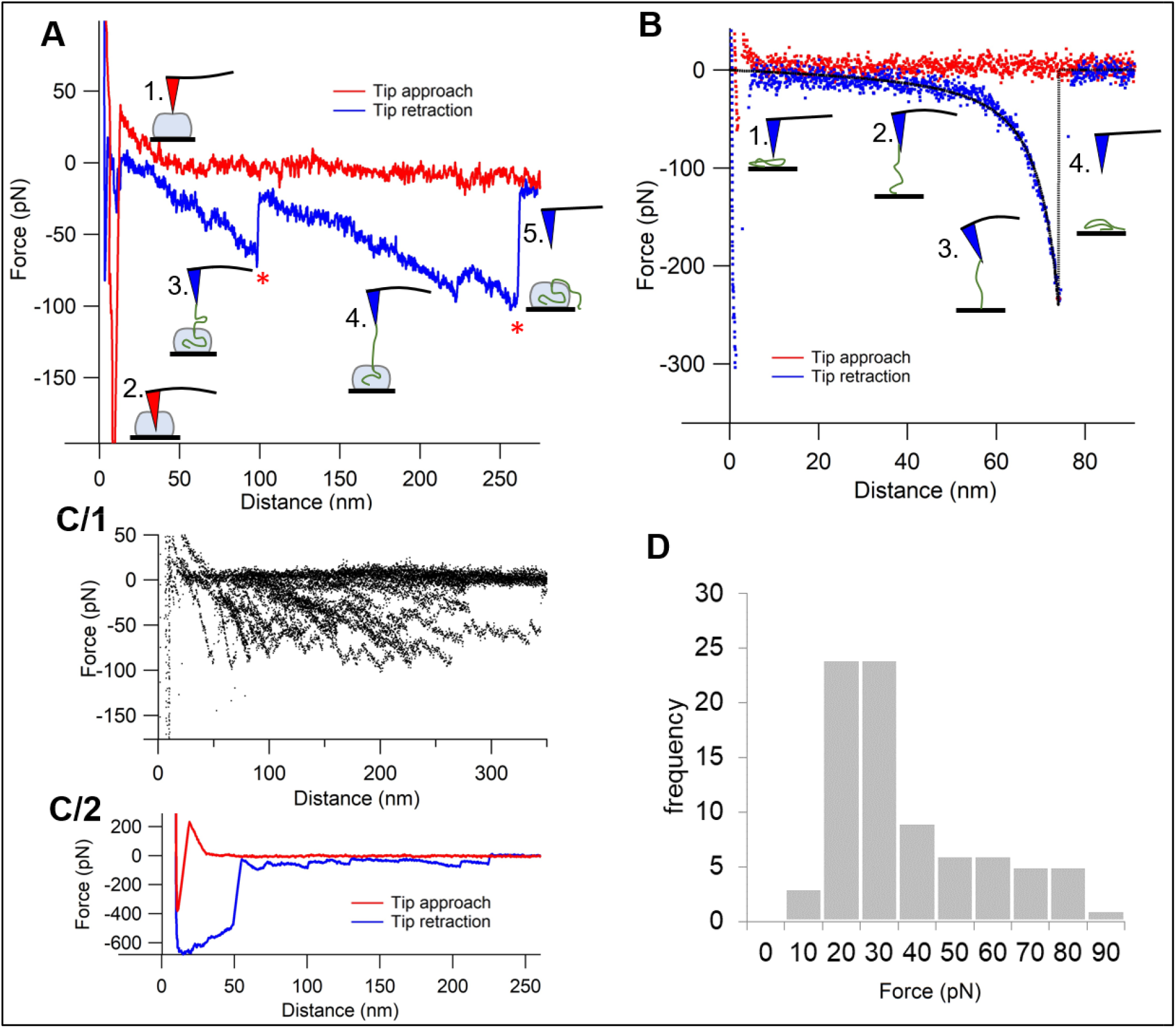
Force spectroscopy of Comirnaty. A) Force-distance curve of a particle indentation (red curve) and RNA extraction (blue curve). The numbered schematics along the curves illustrate the different stages of measurements, with the red and blue tips pointing to the peak “force points”, i.e., the distance where sudden transitions occur. B) Force-distance curve of RNA stretching and its related schematics. Black dashed line shows Worm-like chain model fit (Persistence length = 345 pm, Contour length =83 nm). C/1) Region of interest of RNA extraction force-distance curves superimposed (n=20). C/2) A selected RNA extraction trial with multiple RNA structural transition events. D) Histogram of peak forces detected in RNA extraction curves (peak forces are labeled with red stars in (A).

At a certain distance, a sudden and considerable force drop leading to high negative force values occurred (phase 2) indicating that the particle surface ruptured and resulted in tip attraction. The distance at which rupture took place was ~40% of the contact-point distance (i.e., contact height), suggesting that the particles could be compressed to almost 1/3 of their height during indentation before they lost their mechanical integrity. The force necessary to pierce through the NP surface (~78 pN, Table 2) was 1-3 orders of magnitude lower than that in other ordered nanoscale biomolecular systems, such as liposomes (0.6-1.1 nN).^7, 8^ or empty viral capsids (0.6-5.8 nN).^9–13^

**Table 2.** Nanoparticle biomechanical properties obtained from AFM image and force curve analysis of freshly diluted Comirnaty

| Parameter | Comirnaty |
| --- | --- |
| Vesicle stiffness (pN/nm) | 8.92±8.25 (20) |
| Rupture force (pN) | 77.8±47.2 (20) |
| Attractive force (pN) | 414.4±276.9 (20) |
| Peak force (pN) | 42.7±20.4 (83) |

Entries are Mean ± SD from (n) tests. Further details are in the Methods

Following the negative force regime, a steep rise of force is seen in the force spectra indicating that the tip reached the hard, incompressible glass substrate below the particle. Upon retraction, initially a negative force region is apparent that can be attributed to adhesive forces between the tip and the nanoparticle content. Following this stage, a region containing sawtooth-like transitions appeared in the force trace (marked by red asterisks in Fig. 2A). The steady decrease in force (which corresponds to increase in pulling force) is likely to reflect molecular strands (RNA) being pulled out of the particles, while the sudden force transitions are probably due to either the unfolding of RNA or the rupture of interactions between the RNA and lipid components (phases 3-4). The thread-like behavior reflects the properties of the RNA strand. The superimposition of retraction force traces (Fig. 2C/1) failed to reveal preferred distances at which transitions occur. The number of force sawteeth ranged from a few (as seen in Fig. 2A) to several (Fig. 2C/2). The mean force associated with the sawtooth peaks was 42.7 pN (see peak force distribution in Fig. 2D), which exceeds the force necessary for opening RNA hairpins (13-14 pN),^14^ suggesting that the observed transitions are the result of lipid-RNA interactions rather than RNA unfolding events. At the last transition, during which force returned to zero (i.e., no load on the cantilever), RNA was either pulled completely out of the particle or detached from the AFM tip (phase 5).

In control force spectroscopic measurements collected during indentation cycles performed on the background, force traces corresponding to wormlike chain pulling (WLC) were observed (Fig. 2B). The persistence length calculated from the WLC fits is ~0.3 nm, which is on the same scale as that of ssRNA.^15^ Lack of sawtooth-like transitions indicates no hairpin openings in RNAs being pulled from the supporting surface.

In sum, nanomechanical properties of NPs, especially stiffness, may play an important role in their cell adhesion and internalization, as it was demonstrated earlier for liposomes, extracellular vesicles and viruses.^16–18^ To our knowledge, the present study is the first to analyze a vaccine from this aspect and is unprecedented to show that Comirnaty NPs are rather soft, deformable, highly compliant structures.

### Cryo-TEM of Comirnaty

Consistent with the AFM images of fresh Comirnaty (Fig. 1A) cryo-TEM images of the newly diluted vaccine (Fig. 3 A-C) showed spherical NPs. These NPs exhibit a relatively broad size distribution having about 50 to 200 nm diameter. However, mostly, NPs having 50-80 nm diameters were observed, which may suggest a bimodal distribution. Many of the NPs have a solid core with an electron dense interior, distinct from the diffused aqueous background. This appearance may suggest a winding up of mRNA into “supercoils”. Note that this interior is different from that in the anti-cancer nano-drug Doxil, where a low-density intra-liposome aqueous phase, similar to the background outside the liposome is surrounding the nano-rod-like doxorubicin-sulfate crystal inside the liposome (Fig. 3D).

**Figure 3.**
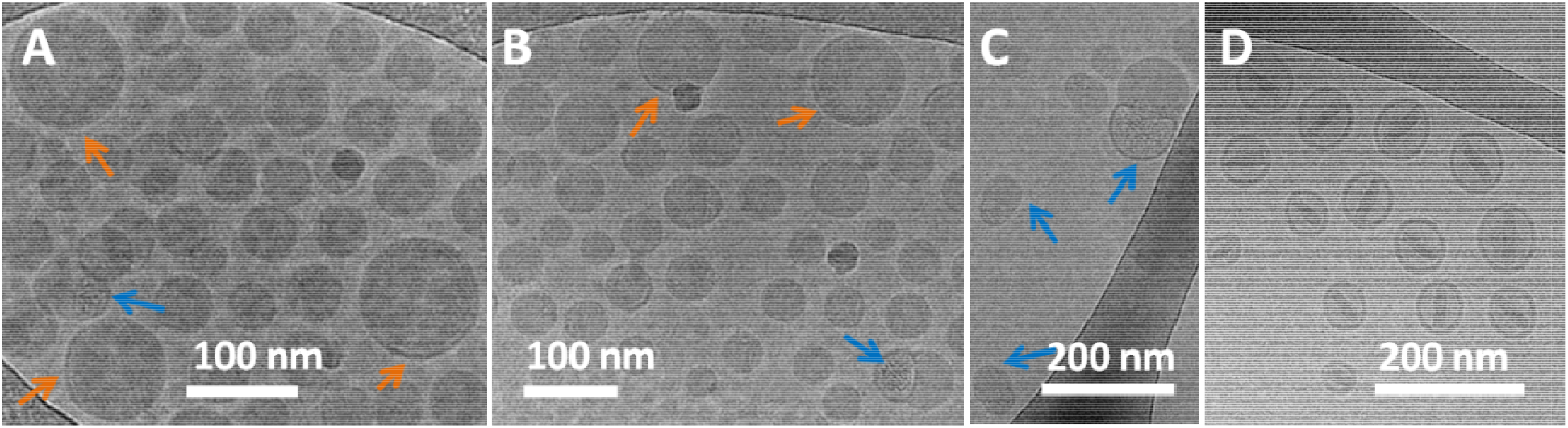
Cryo-TEM images of Comirnaty (A-C) and of Doxil (D). Samples for cryo-TEM (A-C) were processed immediately after thawing the vaccine vial from −80°C and dilution with saline, as instructed by the manufacturer for human application. Orange arrows show the bilayer coating of the NPs. Blue arrows point at bi-compartmental NPs with mRNA containing bleb. D) Cryo-TEM image of Doxil showing the rod-like doxoribicin-sulfate crystal inside the intra-liposome aqueous phase.

The cryo-TEM images of Comirnaty NPs highlight another phenomenon: the presence of electron-dense surface “caps” covering parts of the NPs (orange arrows), which resemble a bilayer coating, as in Doxil (Fig. 3D). Thus, we hypothesize that these thick semicircle lines are phospholipid bilayers, and that Comirnaty NPs are intermittently coated with phospholipid mono- and bi-layers, or there may be membrane-free surface areas possibly covered with polarized water. Additionally, cryo-TEM images of all samples also included particles having a bi-compartmental structure with a bilayer-coated bulge showing “grain-like” nanostructures attributable to the mRNA (blue arrows in Fig. 3 A-C). It was previously suggested that DSPC can be segregated from other lipids forming a bilayer membrane bleb containing aqueous compartments. ^19^

The above observations on intermittent coating of NPs with phospholipid bi- and monolayers of DSPC/PEGylated lipid/cholesterol, and that the NPs may also contain membrane-free surface areas is consistent with the “soft” character of Comirnaty NPs revealed by the AFM measurements.

### Negative staining TEM of Comirnaty enhances the contrast of core structure in stored NPs

The hypothesis on winding up of mRNA in supercoils in Comirnaty LNPs, suggested by the electron-dense grainy appearance of mRNA-LNPs in cryo-TEM images (Figs 3A,B), was further supported by TEM of a sample stored for a week in the refrigerator and then subjected to negative staining with pH 4.5 uranyl acetate. This dried and stained sample (Fig 4) showed a wide variety of odd-shaped LNPs and some molecular details not seen with cryo-TEM were also observed. Higher magnification of some of the globular structures (Fig 4B) shows circular and semicircular chains of ~2-3 nm dots lined up in yarn-ball-like lumps. These may be rationalized as mRNA “supercoils” held tightly together by intra- and interchain forces. Zooming into some structures (red arrows) in Fig. 4A we see intertwined helices winding out from NP remains, probably mRNA-containing fragments (Fig 4C), and other structures seem to capture the process of fusion (Fig. 4A, blue arrows).

**Figure 4.**
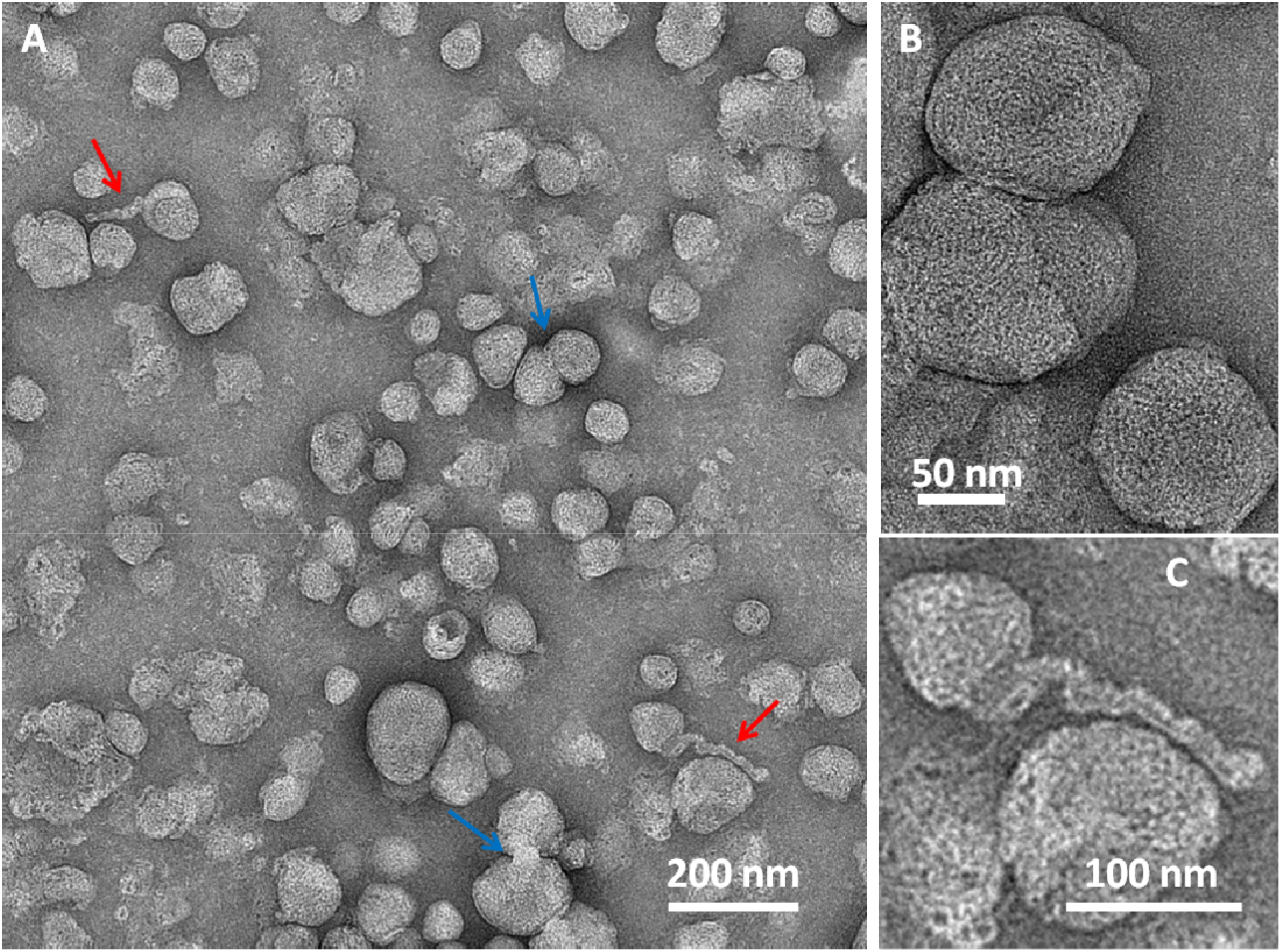
TEM images of Comirnaty sample stored for 7 days in the refrigerator, diluted in water, and stained with acidic uranyl acetate as described in the Method. Blue arrows in (A) point at fusion of liposomes. B) Higher magnification image of some LNPs. C) Zoom into one of the elongated helical-like structures indicated by red arrows in (A).

While the transitional structures could be artifacts in the dried and negatively stained LNPs, breakup and fusion of NPs were also seen in the 1-day stored AFM images (Fig. 1B), and the possibility of helical chains leaving the LNP was also suggested by the stepwise pulling out of string-like structures from Comirnaty by the AFM tip (Fig 2). Furthermore, these transient nanostructures were seen in samples stained with acidic uranyl acetate, and mRNA-LNPs are known to disintegrate in the acidic milieu of endosomes, the site of mRNA release into the cytosol.^20^ Thus, while exposing unforeseen details of LNP core structure, these transient NPs may illustrate the intra-endosomal transformation of LNPs after vaccination, in vivo.

Beyond the acidic milieu, the other likely contributing factor to LNP disintegration is the limited stability of the vaccine in water, as reflected in the brief (up to 6h) shelf-life of Comirnaty on RT after dilution.^42–44^

### Comparing the trans-membrane pH gradient of Comirnaty and Doxil

Transmembrane ion and pH gradients are measures of NP membrane ability to maintain intra-liposome ion (including proton) concentration. Changes and stability of intra-NP proton concentration can be determined for the pH gradient (NP pH ≪ medium pH).^21^ The larger is the pH gradient, the higher is the accumulation of ^14^C-MA in the NPs.^22^ Therefore, we compared the pH gradients in Comirnaty and Doxil, Myocet (non-pegylated liposomal doxorubicin) and Marqibo (non-pegylated liposomal vincristine). In all three liposomal drugs the drug encapsulation is driven by transmembrane pH gradients. In Doxil, this is achieved by the use of transmembrane ammonium sulfate gradient,^23^ while for Myocet^24^ and Marqibo^24^, the pH gradient is ensured by preparing the LNP in citrate buffer, pH 4.0, and raising the pH to neural (pH 7.2) after the active ingredient is encapsulated.^5^. Measurement of the pH gradient in Comirnaty was performed by measuring ^14^C methylamine (^14^C-MA) distribution in Comirnaty and Doxil. This is especially relevant to Comirnaty LNPs as their active ingredient (mRNA) encapsulation is done in a similar exposure to medium pHs, starting at a pH 4.0, followed by changing the medium pH to neutral (pH 7.2).

For all Comirnaty samples, regardless of storage time, there was no ^14^C-MA accumulation by the LNPs, which indicates no pH gradient between the LNP core and the external medium. In contrast, Doxil, which has a similar envelop lipid composition as Comirnaty (DSPC, the main component of HSPC, cholesterol and pegylated lipid DSPC), the measured ΔpH is 1.8 and the calculated internal pH is 4.7.^21^ Similar pH gradients were determined for Myocet and Marqibo.^24^

The loss of internal acidic pH has fundamental impact on the level of ionization of IPC lipid in Comirnaty. ALC 0315 has a pKa of 6.5 and therefore it has a strong cationic character only during LNP formation at pH 4.0.^25, 26^ The rise of intra-LNP pH entails decreased positivity of IPC lipids, and, hence, weakening of their ionic interactions with the mRNA. This could contribute to the gradual disintegration of LNPs after dilution and storage and explains the softness and fragility of Comirnaty NPs after dilution. The increased ion permeability of Comirnaty NPs could be due, at least in part, to the lack of continuous surface bilayer (Fig. 3A,B), since Doxil, which has a similar envelope lipid composition but in the form of continuous bilayer, does not lose the pH gradient in vitro during storage as liposome dispersion and *in vivo*.^23, 27^

### Current LNP models may not be applicable to Comirnaty

There are many features of Comirnaty that cannot be reconciled with the current models of LNP structure developed for small interfering RNAs (siRNAs). In one of these models, referred to as “multilamellar vesicle model”, the 7-8 nm, 21–23 nucleotide-containing siRNA polynucleotides were proposed to be sandwiched between tightly packed IPC enriched monolayers and/or bilayers (Fig. 5A, B).

**Figure 5.**
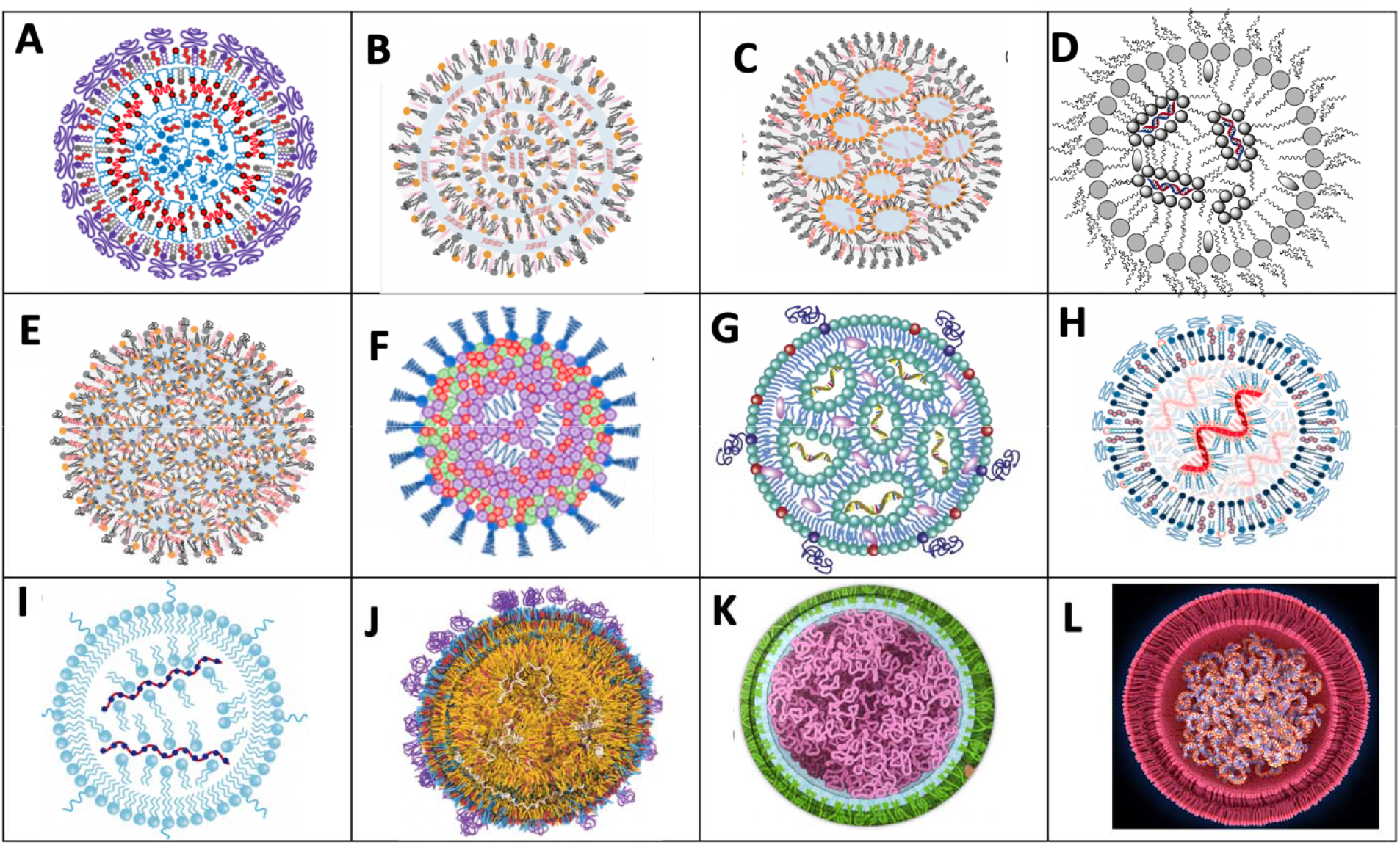
Schematic models of nucleic acid containing LNPs illustrating the variety of concepts. A,B “multilamellar vesicle model” with siRNA sandwiched among the bilayers.^1, 28^ In A) there is an IPC lipid/cholesterol core,^1^ while in B, the lipids are not differentiated.^28^ C) Phospholipid monolayer-covered inverted micelle core.^29^ The siRNA is shown within the surface monolayer and inter-micellar space.^29^ D) Phospholipid-monolayer-coated inverted IPC micelles with siRNA inside the aqueous core of micelles.^30^ E) Phospholipid-monolayer-coated inverted micelles with the siRNA randomly distributed.^29^ F) PEG-lipid coated random assembly of lipids and siRNA.^31^ G) Phospholipid monolayer-coated assembly of inverted IPC lipids, the mRNA is attached to the inner layer of inverted micelles.^32^ H) Phospholipid bilayer coated assembly of mRNA, covered with IPC lipids.^33^ I) Same as H, except the outer membrane is monolayer.^34^ J) mRNA randomly distributed in an amorphous lipid mixture with heterogeneous bilayer surface coat.^35^ K) mRNA thread ball with no identifiable lipid and membrane components. L) Bilayer coated mRNA thread ball with no identifiable lipids.^36^

In another concept, known as “core–shell”, or “nanostructured core” model, the phospholipid monolayer-covered LNPs contain water-filled inverted micelles (Fig. 5C) and the siRNA molecules are in the inter-micellar space and within the outer monolayer membrane. In an alternative of the latter model, the siRNA molecules are located only in the inter-micellar space (Fig. 5D). Fig 5E and F show further models wherein the relative positions of lipids and siRNA are vague.

As for mRNA LNPs, such as Comirnaty, one must consider that the ~1,414 kDa, 4,284 nucleotide-containing mRNA in Comirnaty has an extended length of ~1,500 nm, which is roughly 180-times longer than a siRNA. The huge size difference between siRNA and mRNA has been well recognized,^37^ yet this dissimilarity is not illustrated in most schematic cartoons of mRNA-LNP models. For example, in one, the minimally larger mRNA (compared to siRNA) is shown as being bound to the charged inner layer of inverted micelles made of IPC (Fig. 5G), or as spirals covered by IPC lipids (Fig. 5H, I). Yet another model of mRNA-LNP shows randomly distributed mRNA in an amorphous lipid core. The envelope of mRNA-LNPs is variably shown as phospholipid enriched monolayer (Fig. 5G, I) or bilayer (Fig. 5H).

Taken together, visual representations of mRNA-LNP models are still far from being to scale, the lamellarity of the outer membrane is ambiguous, and the relationship between the mRNA and lipids is highly variable in the different models. These facts highlight the need to better understand the molecular structure of mRNA-LNPs and, accordingly, better visualize them in schematic model.

### Proposal of a new mRNA vaccine specific LNP model

According to Buschmann et al., each Comirnaty LNP contains up to 10 mRNA molecules.^28^ Considering the above-mentioned size of Comirnaty mRNA, they cannot fit into 60-150 nm diameter spheres unless they tightly wind up in dense loops or supercoils, just as the DNA fills the nuclei. There exist such vaccine presentations in the public media, showing the mRNA in the vaccine LNPs as balls of yarn, or thread balls (Fig. 5K, L), but these simplified, fictional models make no attempt to illustrate the details of the membranes and the relationships among the different internal components, most critically the IPC lipids, whose number exceeds that of the mRNA nucleotides 6-fold.^38^

The analytic and visual data in the present study, taken together with the mentioned size information suggest a new model wherein a single, or multiple copies of mRNA-IPC complexes densely fill up part, or the whole internal space of LNPs, and the IPC lipids, or self-associated IPC clusters, stabilize the tertiary structure of mRNA supercoils via intra- and interloop crosslinks and bridges (Fig. 6A). Crosslinks may be provided by hydrogen bonds between the free =O and -OH groups of IPC lipids and the nitrogenous bases in the mRNA, while bridges may be formed from IPC clusters.^6^

**Figure 6.**
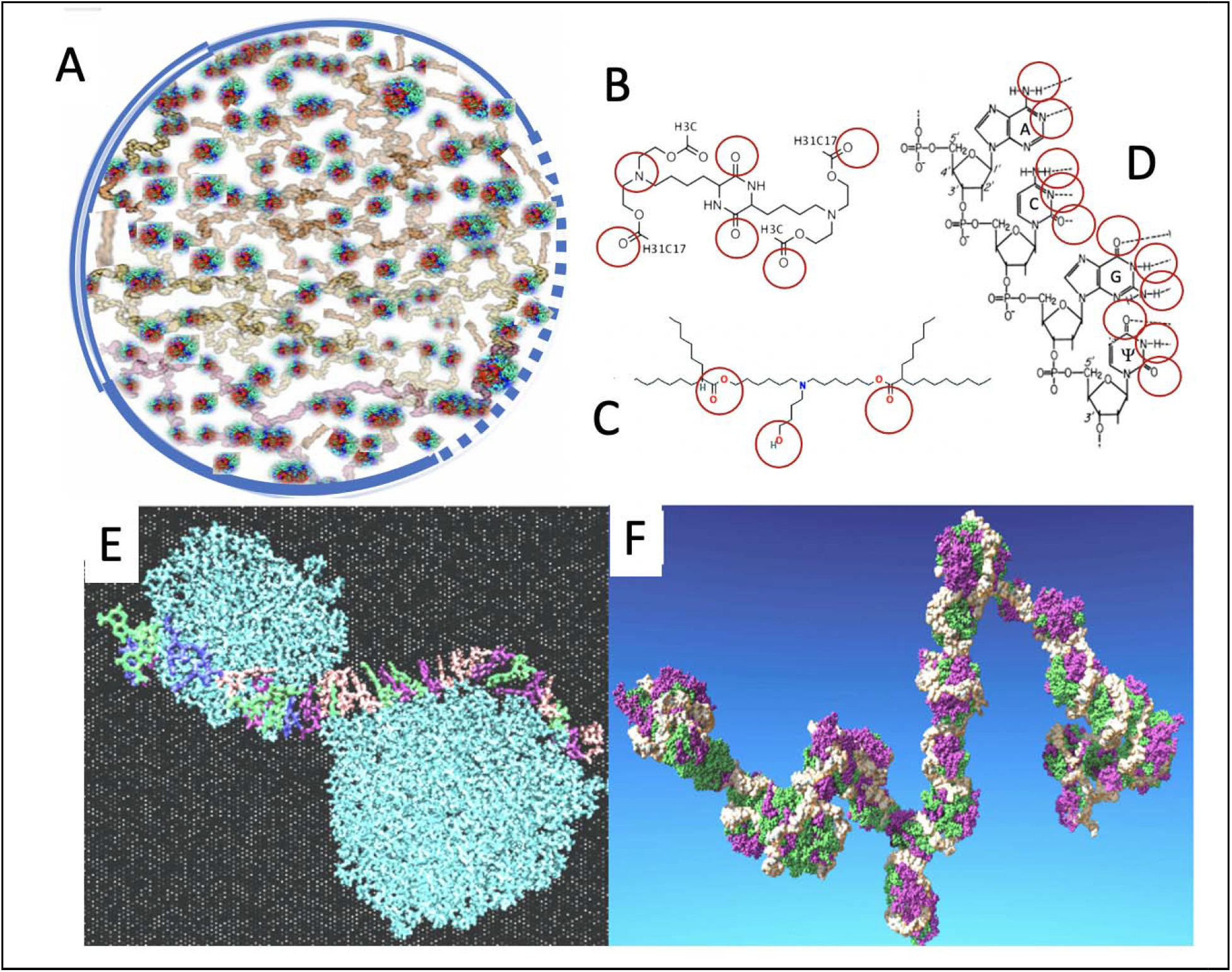
A) Scheme of a new Comirnaty-specific LNP model wherein the stability of yarn ball-like mRNA lumps (gray) is secured by IPC lipid crosslinks and clusters (green and red dots). The NP is surrounded by phospholipid bilayer (double line), monolayer (single line) or no membrane (dotted line). B) Chemical structure of DML, containing a diketopiperazine core, two methyl ester end-groups and two linoleic ethyl ester end-groups.^6^ C) Chemical structure of ALC-0315, the ionizable cationic lipid in Comirnaty [(4-hydroxybutyl) azanediyl)bis (hexane-6,1-diyl)bis(2-hexyldecanoate (Table S1 and Ref. ^38^); D) A four nucleotide section of the mRNA chain in Comirnaty showing the N and O atoms available for hydrogen bonding, ψ, pseudouridine; E) The Rissanau et. al model of DML lipid cluster-mRNA complexation showing a 30 nucleotide-containing short mRNA meandering along 2 DML clusters. Reproduced from Ref. 6 with permission. F) Molecular dynamics simulation of a 642-nucleotide-containing mRNA complexed with multiple DML clusters that protect the mRNA from hydrolysis. Reproduced from Ref. 6 with permission.

The stability of supercoils may be secured in the frozen vaccine by the fact that the molecules do not vibrate as vigorously as in unfrozen state,^39^ and it may be increased by electrostatic interactions with IPC lipids as a function of their positivity. However, both the hydrogen bonds, which are considered as weak bonds, and the electrostatic bonds, whatever still exists at neutral pH, become weaker as the temperature increases, as occurs during storage of the aqueous dispersion on RT, and even faster, when the mRNA-LNP reaches the temperature of body fluids (37°C).

The above weakly stabilized lipoplex model of Comirnaty mRNA-LNP was inspired by a recent study by Rissanou et al.^6^ showing that an IPC lipid (called DML, details are seen in the legend of Fig. 6), which is very similar to Comirnaty’s IPC (ALC 0315) in terms of molecular size and steric positions of free OH- and O= groups, which can theoretically align with similar groups on the purine and pyrimidine bases of single stranded mRNA (red circles in Fig. 6B).

For DML, evidence was provided that it can self-associate into clusters that can bind to mRNA both by ionic and hydrogen bonds,^6^ and thus stabilize the tertiary structure of the complex in an unprecedented molecular assembly where the mRNA meanders around lipid clusters (Fig. 6E and F). We hypothesize that ALC 0315 in Comirnaty may have similar mRNA binding capability and stabilize Comirnaty’s mRNA in the delineated supercoil conformation, in a kind of glomerular lattice.

## Outlook

This study provided evidence that the NPs in Comirnaty are not conventional bilayer-coated liposomes, nor are identical to siRNA-carrying LNPs, but represent a unique molecular assembly whose structure can most easily be imagined as mRNA supercoils stabilized by pH and temperature-dependent weak inter- and intramolecular interactions, involving hydrogen and, possibly, ionic bonds. These NPs are “softer” than the standard bilayer liposomes or other LNPs with fuller membrane coverage, which may explain the diluted vaccine’s limited stability on ambient temperature.^42–44^

Despite its softness, and, hence, limited stability, the vaccine is clearly very effective in inducing an immune response, raising the possibility that the above newly recognized structural features are contributing to, or may be essential for its immunogenic function. Our data may therefore lay the grounds for a paradigm shift in understanding the structure-function relationship in mRNA vaccines, as well as contribute to developing better imaging of LNP core structure.

## Materials and Methods

### Materials

BNT162b2 (Comirnaty) was from Pfizer/BioNtech, the vaccine used for human vaccinations against SARS-CoV-2 infections. The frozen vials were thawed and then diluted with saline by a healthcare professional, as instructed by the manufacturer. Doxil was from a local pharmacy. The composition of the vaccine and Doxil are tabulated in Supplementary Table S1.

### AFM imaging and force spectroscopy

For AFM experiments, unused, sterile batch of Comirnaty vaccine, just prior to the end of its warranty period, was diluted fresh in physiological salt solution. The diluted sample was used immediately. Further experiments were done with samples (a) stored at 4°C for one day or (b) frozen in liquid N_2_, stored at −20°C for one day and thawed to room temperature. Round coverslips (Ted Pella Inc., Redding, CA) were glued with UV epoxy onto metal AFM specimen disks, and cleaned with ethanol, then methanol. The cleaned glass surface was dried in nitrogen stream. 100 μl vaccine was dropped onto the freshly cleaned surface and incubated for 20 min at room temperature. Then the surface was washed 5 times with 100 μl PBS (16 mM Na_2_PO_4_, 4 mM NaHPO_4_, 150 mM NaCl; pH 7.4). Imaging was carried out in tapping mode at 25°C with a Cypher ES AFM (Asylum Research, Santa Barbara, CA) using BL-AC40TS cantilever (silicon nitride, nominal stiffness: 90 pN/nm, tip radius: 8 nm; Olympus, Japan) at typical line rates of 0.5 Hz. The cantilever was oscillated photothermally (BlueDrive) at its resonance frequency (typically 20 kHz in water). Prior to the measurements, the cantilevers were calibrated by using the thermal method in air.^40^

In situ force spectroscopy was carried out in contact mode on vesicles selected in a previously scanned AFM image. During force spectroscopy, the cantilever was moved vertically with a speed of 1 μm/s from a height of 500 nm towards the vesicle vertex until a force threshold of 5 nN was reached. Then the tip was immediately retracted with the same speed. Deflection of the cantilever, hence force as a function of cantilever position (force-indentation curve or force curve), was recorded during the process. Images and force spectroscopy data were analyzed by using the-built in algorithms of the AFM driving software (IgorPro, WaveMetrics Inc., Lake Oswego, OR).

### Measurement of Comirnaty NP size and size distribution by dynamic light scattering

Dynamic light scattering (DLS) measurements were performed immediately after thawing a Comirnaty vial and diluting NPs with saline (performed by a healthcare professional). Size (diameter, D), size distribution (polydispersity index, PDI and SPAN) and zeta potential (Zp) of the NPs were determined with a Malvern Zetasizer Nano ZS instrument (Malvern, Worcestershire, UK) at an angle of 173°. In addition to the PDI, a standard measure of homogeneity, we also determined the SPAN, a better measure of distribution broadness than PDI. It is defined as D_90_–D_10_/D_50_ where D_10, 50 and 90_ are the percentiles under the size distribution curve.

### TEM and Cryo-TEM imaging

Negative-staining transmission electron microscopy (TEM) was performed by applying a drop (3 μl) of sample to a glow-discharged TEM grid (carbon supported film on 300 mesh Cu grids, Ted Pella, Ltd). After 30 sec the excess liquid was blotted, the grids were stained with 2% uranyl acetate for 30 sec and allowed to dry in air. Imaging was carried out using a FEI Tecnai 12 G2 Twin TEM operated at 120 kV. The images were recorded by a 4Kx4K FEI Eagle CCD camera using the TIA software. Direct imaging of samples by cryogenic transmission electron microscopy (cryo-TEM) was performed as described elsewhere.^27^ Three μL of sample was applied onto a glow-discharged 300 mesh copper TEM grid coated with a holey carbon film (Lacey substrate Ted Pella, Ltd.). The excess liquid was blotted and the specimens were vitrified by rapid plunging into liquid ethane pre-cooled by liquid nitrogen using Vitrobot Mark IV (FEI). We then transferred the vitrified samples into a cryo specimen holder (Gatan model 626; Gatan Inc.) and imaged them at −177°C using a Tecnai 12 G2 Twin TEM (FEI), operated at an acceleration voltage of 120 kV in low-dose mode. Images were recorded with a 4Kx4K FEI Eagle CCD camera. TIA (Tecnai Imaging & Analysis) software was used to record the images.

### Measurement of intra-LNP pH/pH gradient

The assay, originally described by Abraham et al.,^22^ consisted of measuring the distribution of radioactive methylamine (^14^C-MA) between the NPs and solvent medium, as applied earlier for Doxil.^21^ In brief, ^14^C-MA was added to freshly diluted Comirnaty LNPs. After 20 min incubation at 55 °C, the LNP dispersion was divided equally into two parts. One part of the LNP solution was ultracentrifuged using Amicon Ultra-15 tubes using a filter of 100 K molecular weight cutoff (Millipore, MA) for the separation of the unencapsulated (extra-LNP) from the encapsulated ^14^C-MA. The medium collected after ultracentrifugation and the other part of the original liposome dispersion (before centrifugation) were first treated with Opti-Fluor (Perkin Elmer, Waltham, MA) and stored overnight at 2–8 °C before scintillation counting of the radioactivity by a β-counter. The ratio of ^14^C-MA between the LNPs and the extra-LNP medium was used to calculate the transmembrane pH gradient as follows: ΔpH = Log{[H^+^]inside/[H^+^]outside = Log{[methylamine] inside/[methylamine] outside.

## Data availability

Source data are provided with this paper and its Supplementary STable 1. All other data supporting the findings of this study are available from the corresponding author upon reasonable request.

## Author contributions

JS, MK and YB conceived the idea and designed the experiments, K.T, T.B, Y.L.K and B.K. performed the measurements and analyzed the data, and the paper was written with contributions from all authors. All authors agreed with the manuscript.

## SUPPLEMENT

**Table S1.**
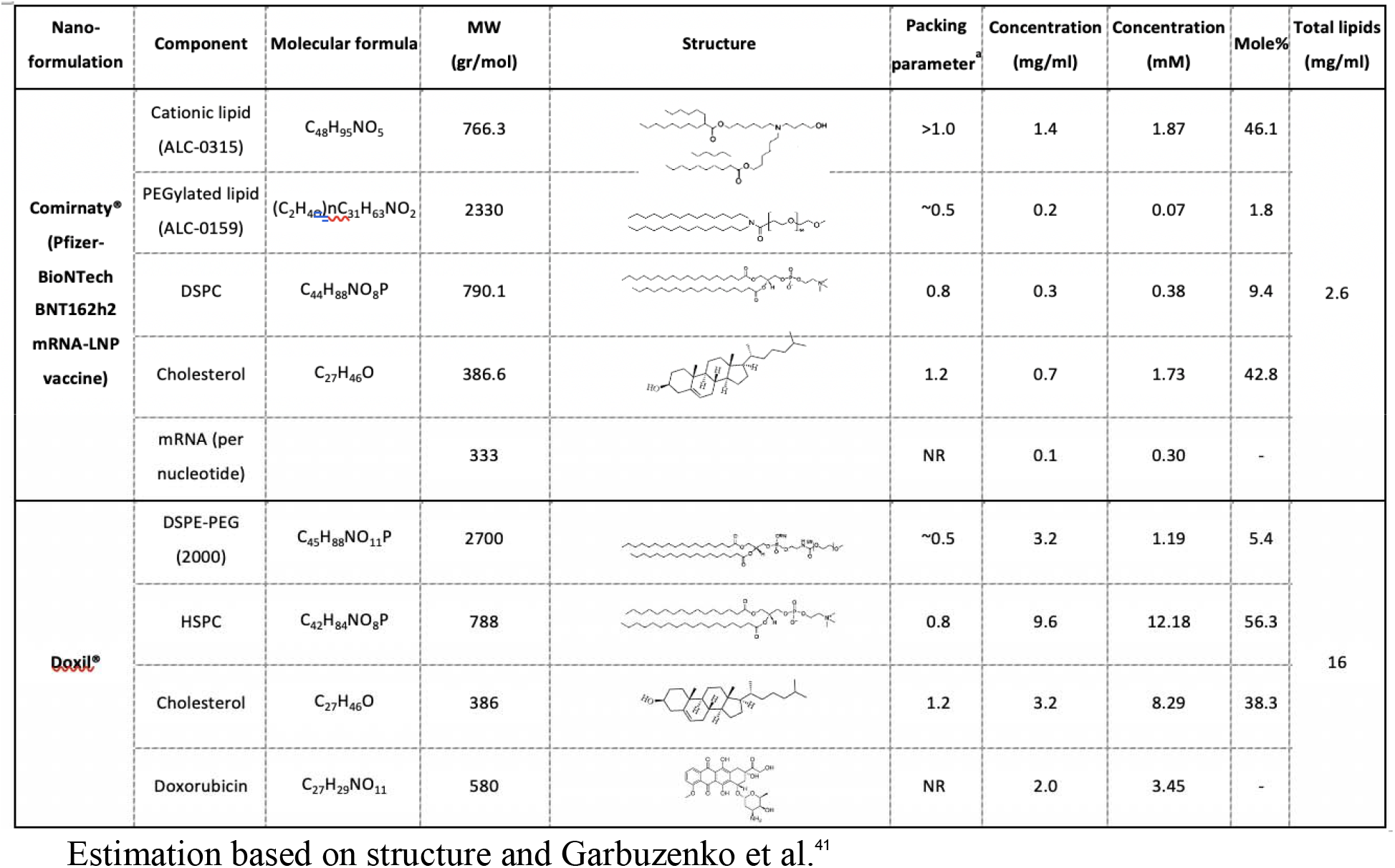
Composition and other characteristics of Comirnaty and Doxil

## References

1. Kulkarni, J. A.; Darjuan, M. M.; Mercer, J. E.; Chen, S.; van der Meel, R.; Thewalt, J. L.; Tam, Y. Y. C.; Cullis, P. R., On the Formation and Morphology of Lipid Nanoparticles Containing Ionizable Cationic Lipids and siRNA. ACS Nano 2018, 12 (5), 4787–4795.

2. Witzigmann, D.; Kulkarni, J. A.; Leung, J.; Chen, S.; Cullis, P. R.; van der Meel, R., Lipid nanoparticle technology for therapeutic gene regulation in the liver. Adv Drug Deliv Rev 2020, 159, 344–363.

3. Hafez, I. M.; Maurer, N.; Cullis, P. R., On the mechanism whereby cationic lipids promote intracellular delivery of polynucleic acids. Gene Ther 2001, 8 (15), 1188–96.

4. Allen, T. M.; Cullis, P. R., Drug delivery systems: entering the mainstream. Science 2004, 303 (5665), 1818–22.

5. Allen, T. M.; Cullis, P. R., Liposomal drug delivery systems: from concept to clinical applications. Adv Drug Deliv Rev 2013, 65 (1), 36–48.

6. Rissanou, A. N.; Ouranidis, A.; Karatasos, K., Complexation of single stranded RNA with an ionizable lipid: an all-atom molecular dynamics simulation study. Soft Matter 2020, 16 (30), 6993–7005.

7. Delorme, N.; Fery, A., Direct method to study membrane rigidity of small vesicles based on atomic force microscope force spectroscopy. Phys Rev E Stat Nonlin Soft Matter Phys 2006, 74 (3 Pt 1), 030901.

8. Li, S.; Eghiaian, F.; Sieben, C.; Herrmann, A.; Schaap, I. A. T., Bending and puncturing the influenza lipid envelope. Biophys J 2011, 100 (3), 637–645.

9. Roos, W. H.; Gertsman, I.; May, E. R.; Brooks, C. L., 3rd; Johnson, J. E.; Wuite, G. J., Mechanics of bacteriophage maturation. Proc Natl Acad Sci U S A 2012, 109 (7), 2342–7.

10. Snijder, J.; Reddy, V. S.; May, E. R.; Roos, W. H.; Nemerow, G. R.; Wuite, G. J., Integrin and defensin modulate the mechanical properties of adenovirus. J Virol 2013, 87 (5), 2756–66.

11. Roos, W. H.; Radtke, K.; Kniesmeijer, E.; Geertsema, H.; Sodeik, B.; Wuite, G. J., Scaffold expulsion and genome packaging trigger stabilization of herpes simplex virus capsids. Proc Natl Acad Sci U S A 2009, 106 (24), 9673–8.

12. Ivanovska, I.; Wuite, G.; Jonsson, B.; Evilevitch, A., Internal DNA pressure modifies stability of WT phage. Proc Natl Acad Sci U S A 2007, 104 (23), 9603–8.

13. Michel, J. P.; Ivanovska, I. L.; Gibbons, M. M.; Klug, W. S.; Knobler, C. M.; Wuite, G. J.; Schmidt, C. F., Nanoindentation studies of full and empty viral capsids and the effects of capsid protein mutations on elasticity and strength. Proc Natl Acad Sci U S A 2006, 103 (16), 6184–9.

14. Bercy, M.; Bockelmann, U., Hairpins under tension: RNA versus DNA. Nucleic Acids Res 2015, 43 (20), 9928–36.

15. Hayashi, K.; Chaya, H.; Fukushima, S.; Watanabe, S.; Takemoto, H.; Osada, K.; Nishiyama, N.; Miyata, K.; Kataoka, K., Influence of RNA Strand Rigidity on Polyion Complex Formation with Block Catiomers. Macromol Rapid Commun 2016, 37 (6), 486–93.

16. Sakai-Kato, K.; Yoshida, K.; Takechi-Haraya, Y.; Izutsu, K. I., Physicochemical Characterization of Liposomes That Mimic the Lipid Composition of Exosomes for Effective Intracellular Trafficking. Langmuir 2020, 36 (42), 12735–12744.

17. Hui, Y.; Yi, X.; Wibowo, D.; Yang, G.; Middelberg, A. P. J.; Gao, H.; Zhao, C. X., Nanoparticle elasticity regulates phagocytosis and cancer cell uptake. Sci Adv 2020, 6 (16), eaaz4316.

18. Kiss, B.; Mudra, D.; Torok, G.; Martonfalvi, Z.; Csik, G.; Herenyi, L.; Kellermayer, M., Single-particle virology. Biophys Rev 2020, 12 (5), 1141–1154.

19. Leung, A. K.; Tam, Y. Y.; Chen, S.; Hafez, I. M.; Cullis, P. R., Microfluidic Mixing: A General Method for Encapsulating Macromolecules in Lipid Nanoparticle Systems. J Phys Chem B 2015, 119 (28), 8698–706.

20. Hald Albertsen, C.; Kulkarni, J. A.; Witzigmann, D.; Lind, M.; Petersson, K.; Simonsen, J. B., The role of lipid components in lipid nanoparticles for vaccines and gene therapy. Adv Drug Deliv Rev 2022, 188, 114416.

21. Wei, X.; Shamrakov, D.; Nudelman, S.; Peretz-Damari, S.; Nativ-Roth, E.; Regev, O.; Barenholz, Y., Cardinal Role of Intraliposome Doxorubicin-Sulfate Nanorod Crystal in Doxil Properties and Performance. ACS Omega 2018, 3 (3), 2508–2517.

22. Abraham, S. A.; Edwards, K.; Karlsson, G.; MacIntosh, S.; Mayer, L. D.; McKenzie, C.; Bally, M. B., Formation of transition metal-doxorubicin complexes inside liposomes. Biochim Biophys Acta 2002, 1565 (1), 41–54.

23. Barenholz, Y., Doxil(R)--the first FDA-approved nano-drug: lessons learned. J Control Release 2012, 160 (2), 117–34.

24. Madden, T. D.; Harrigan, P. R.; Tai, L. C.; Bally, M. B.; Mayer, L. D.; Redelmeier, T. E.; Loughrey, H. C.; Tilcock, C. P.; Reinish, L. W.; Cullis, P. R., The accumulation of drugs within large unilamellar vesicles exhibiting a proton gradient: a survey. Chem Phys Lipids 1990, 53 (1), 37–46.

25. Semple, S. C.; Akinc, A.; Chen, J.; Sandhu, A. P.; Mui, B. L.; Cho, C. K.; Sah, D. W.; Stebbing, D.; Crosley, E. J.; Yaworski, E.; Hafez, I. M.; Dorkin, J. R.; Qin, J.; Lam, K.; Rajeev, K. G.; Wong, K. F.; Jeffs, L. B.; Nechev, L.; Eisenhardt, M. L.; Jayaraman, M.; Kazem, M.; Maier, M. A.; Srinivasulu, M.; Weinstein, M. J.; Chen, Q.; Alvarez, R.; Barros, S. A.; De, S.; Klimuk, S. K.; Borland, T.; Kosovrasti, V.; Cantley, W. L.; Tam, Y. K.; Manoharan, M.; Ciufolini, M. A.; Tracy, M. A.; de Fougerolles, A.; MacLachlan, I.; Cullis, P. R.; Madden, T. D.; Hope, M. J., Rational design of cationic lipids for siRNA delivery. Nat Biotechnol 2010, 28 (2), 172–6.

26. Hajj, K. A.; Ball, R. L.; Deluty, S. B.; Singh, S. R.; Strelkova, D.; Knapp, C. M.; Whitehead, K. A., Branched-Tail Lipid Nanoparticles Potently Deliver mRNA In Vivo due to Enhanced Ionization at Endosomal pH. Small 2019, 15 (6), e1805097.

27. Nordstrom, R.; Zhu, L.; Harmark, J.; Levi-Kalisman, Y.; Koren, E.; Barenholz, Y.; Levinton, G.; Shamrakov, D., Quantitative Cryo-TEM Reveals New Structural Details of Doxil-Like PEGylated Liposomal Doxorubicin Formulation. Pharmaceutics 2021, 13 (1).

28. Buschmann, M. D.; Carrasco, M. J.; Alishetty, S.; Paige, M.; Alameh, M. G.; Weissman, D., Nanomaterial Delivery Systems for mRNA Vaccines. Vaccines (Basel) 2021, 9 (1).

29. Viger-Gravel, J.; Schantz, A.; Pinon, A. C.; Rossini, A. J.; Schantz, S.; Emsley, L., Structure of Lipid Nanoparticles Containing siRNA or mRNA by Dynamic Nuclear Polarization-Enhanced NMR Spectroscopy. J Phys Chem B 2018, 122 (7), 2073–2081.

30. Zatsepin, T. S.; Kotelevtsev, Y. V.; Koteliansky, V., Lipid nanoparticles for targeted siRNA delivery - going from bench to bedside. Int J Nanomedicine 2016, 11, 3077–86.

31. Sebastiani, F.; Yanez Arteta, M.; Lerche, M.; Porcar, L.; Lang, C.; Bragg, R. A.; Elmore, C. S.; Krishnamurthy, V. R.; Russell, R. A.; Darwish, T.; Pichler, H.; Waldie, S.; Moulin, M.; Haertlein, M.; Forsyth, V. T.; Lindfors, L.; Cardenas, M., Apolipoprotein E Binding Drives Structural and Compositional Rearrangement of mRNA-Containing Lipid Nanoparticles. ACS Nano 2021, 15 (4), 6709–6722.

32. Forchette, L.; Sebastian, W.; Liu, T., A Comprehensive Review of COVID-19 Virology, Vaccines, Variants, and Therapeutics. Curr Med Sci 2021, 41 (6), 1037–1051.

33. Nanotechnology, N. C. f. S., Sustainable Nano. a blog by the NSF Center for Sustainable Nanotechnology 2022, https://sustainable-nano.com/2021/12/02/lipid-nanoparticles-covid-vaccines/.

34. Heinz, F. X.; Stiasny, K., Distinguishing features of current COVID-19 vaccines: knowns and unknowns of antigen presentation and modes of action. NPJ Vaccines 2021, 6 (1), 104.

35. Tenchov, R.; Bird, R.; Curtze, A. E.; Zhou, Q., Lipid Nanoparticles-From Liposomes to mRNA Vaccine Delivery, a Landscape of Research Diversity and Advancement. ACS Nano 2021.

36. Scientist, N., The mRNA technology behind covid-19 vaccines can transform medicine. New Scientist 2021 Oct 13, https://www.newscientist.com/article/mg25133562-800-the-mrna-technology-behind-covid-19-vaccines-can-transform-medicine/.

37. Schoenmaker, L.; Witzigmann, D.; Kulkarni, J. A.; Verbeke, R.; Kersten, G.; Jiskoot, W.; Crommelin, D. J. A., mRNA-lipid nanoparticle COVID-19 vaccines: Structure and stability. Int J Pharm 2021, 601, 120586.

38. Szebeni, J.; Storm, G.; Ljubimova, J. Y.; Castells, M.; Phillips, E. J.; Turjeman, K.; Barenholz, Y.; Crommelin, D. J. A.; Dobrovolskaia, M. A., Applying lessons learned from nanomedicines to understand rare hypersensitivity reactions to mRNA-based SARS-CoV-2 vaccines. Nat Nanotechnol 2022, 17 (4), 337–346.

39. Meot-ner, M.; Sieck, L. W. J. J. o. t. A. C. S., The ionic hydrogen bond. 1. Sterically hindered bonds. Solvation and clustering of protonated amines and pyridines. 1983, 105, 2956–2961.

40. Hutter, J. L.; Bechhoefer, J., Calibration of atomic□force microscope tips. Review of Scientific Instruments 1993, 64, 1868–73.

41. Garbuzenko, O.; Zalipsky, S.; Qazen, M.; Barenholz, Y., Electrostatics of PEGylated micelles and liposomes containing charged and neutral lipopolymers. Langmuir 2005, 21, 2560–2568.

42. Pfizer/BioNTech, COMIRNATY^®^ (COVID-19 Vaccine, mRNA) suspension for injection, for intramuscular use. https://www.fda.gov/media/151707/download 2022.

43. Crommelin, D. J. A.; Anchordoquy, T. J.; Volkin, D. B.; Jiskoot, W.; Mastrobattista, E., Addressing the Cold Reality of mRNA Vaccine Stability. J Pharm Sci 2021, 110 (3), 997–1001.

44. Blenke, E. O.; Ornskov, E.; Schoneich, C.; Nilsson, G.; Volkin, D. B.; Mastrobattista, E.; Almarsson, O.; Crommelin, D. J. A., The storage and in-use stability of mRNA vaccines and therapeutics: Not a cold case. J Pharm Sci 2022.

